# SARS-CoV-2-host chimeric RNA-sequencing reads do not necessarily signify virus integration into the host DNA

**DOI:** 10.1101/2021.03.05.434119

**Authors:** Anastasiya Kazachenka, George Kassiotis

**Affiliations:** Retroviral Immunology, The Francis Crick Institute, 1 Midland Road, London NW1 1AT, UK; Department of Infectious Disease, St Mary’s Hospital, Imperial College London, London W2 1PG, UK

## Abstract

The human genome bears evidence of extensive invasion by retroviruses and other retroelements, as well as by diverse RNA and DNA viruses. High frequency of somatic integration of the RNA virus severe acute respiratory syndrome coronavirus 2 (SARS-CoV-2) into the DNA of infected cells was recently suggested, partly based on the detection of chimeric RNA-sequencing (RNA-seq) reads between SARS-CoV-2 RNA and RNA transcribed from human host DNA. Here, we examined the possible origin of human-SARS-CoV-2 chimeric reads in RNA-seq libraries and provide alternative explanations for their origin. Chimeric reads were frequently detected also between SARS-CoV-2 RNA and RNA transcribed from mitochondrial DNA or episomal adenoviral DNA present in transfected cell lines, which was unlikely the result of SARS-CoV-2 integration. Furthermore, chimeric reads between SARS-CoV-2 RNA and RNA transcribed from nuclear DNA was highly enriched for host exonic, than intronic or intergenic sequences and often involved the same, highly expressed host genes. These findings suggest that human-SARS-CoV-2 chimeric reads found in RNA-seq data may arise during library preparation and do not necessarily signify SARS-CoV-2 reverse transcription, integration in to host DNA and further transcription.

## INTRODUCTION

Viruses hijack the host cell to replicate their RNA or DNA genomes and create progeny virions. An extreme form of viral parasitism is the integration of a viral genome DNA copy into the host cell DNA (Burns and Boeke, 2012;Feschotte and Gilbert, 2012). Although diverse classes of RNA viruses create a complementary DNA (cDNA) copy through reverse-transcription of their genomes during their life cycle, integration into the host DNA is a characteristic obligatory step for retroviruses, as well as for endogenous retroelements (Coffin et al., 1997;Burns and Boeke, 2012;Feschotte and Gilbert, 2012).

The machinery that mediates reverse transcription and integration of the retroviral and endogenous retroelement genomes can also use alternative RNA templates, creating genomic cDNA copies of the latter. For example, mammalian apparent long terminal repeat (LTR)-retrotransposons (MaLRs) rely on endogenous retroviruses (ERVs) for their reverse-transcription and integration. Similarly, short interspersed nuclear elements (SINEs), including *Alu* elements, rely on long interspersed nuclear elements (LINEs) for their reverse transcription and integration (Coffin et al., 1997;Burns and Boeke, 2012;Feschotte and Gilbert, 2012).

The reverse transcriptase and endonuclease activity of LINEs, carried out by the ORF2p protein, can also mediate reverse transcription and integration of unrelated viral and non-viral RNAs (Klenerman et al., 1997;Esnault et al., 2000;Buzdin, 2004). Indeed, the human genome contains DNA copies of distinct RNA and DNA viruses (Blinov et al., 2017), as well as numerous retrogenes and pseudogenes (Baertsch et al., 2008;Richardson et al., 2014;Staszak and Makałowska, 2021), highlighting the possible, albeit infrequent, reverse transcription and integration of non-retroviral RNA into the host genome.

Recent studies reported a high frequency of reverse transcription and integration of severe acute respiratory syndrome coronavirus 2 (SARS-CoV-2) RNA in infected cells (Zhang et al., 2020;Ying et al., 2021), with implications for diagnostic detection of SARS-CoV-2 nucleic acids by RT-qPCR and for viral antigen persistence. These findings were partly based on the identification of chimeric reads between viral and human RNA in next-generation RNA-sequencing (RNA-seq) data (Zhang et al., 2020;Ying et al., 2021). Here, we examined the potential source of such chimeric reads and found that they are more likely to be a methodological product, than the result of genuine reverse transcription, integration and expression.

## METHODS

### RNA-seq analysis

Public RNA-seq datasets (Blanco-Melo et al., 2020) under the accession number GSE147507 were downloaded from NCBI Gene Expression Omnibus (GEO) server. Adapter and quality trimming were conducted using Trimmomatic v0.36 (Bolger et al., 2014). Quality of sequencing reads was assessed by FastQC v0.11.5. The resulted reads were aligned to the merged GRCh38/hg38 genome (including alternative and random chromosome sequences) and SARS-CoV-2 NC_045512v2 genome using STAR v2.7.1 aligner (Dobin et al., 2013). GENCODE v29 basic version and wihCor1 NCBI genes (http://hgdownload.soe.ucsc.edu/goldenPath/wuhCor1/bigZips/genes/) were used for human and SARS-CoV-2 gene annotations respectively. Chimeric reads were called using STAR parameters as used in prior reports (Zhang et al., 2020). Gene expression was calculated by FeatureCounts (part of the Subread package v1.5.0) (Liao et al., 2014) and normalized with DESeq2 v1.22.1 within R v3.5.1 (Love et al., 2014). The Integrative Genomics Viewer (IGV) v2.5.3 was used to visualize aligned non-chimeric and chimeric reads (Robinson et al., 2011). BLASTN+ v2.3.0 was used to align mtRNA-nRNA chimeric reads to identify mitochondrial and nuclear aligning sequences within the reads (Camacho et al., 2009). Reads containing highly homologous sequences to mitochondrial and nuclear genomes simultaneously were removed from analysis. Viral-host chimeric reads were aligned to SARS-CoV-2 and human reference genomes using the same method to quantify overlapping regions between viral and human genome aligning parts of the reads.

## RESULTS

### Human-SARS-CoV-2 chimeric reads in RNA-seq libraries of SARS-CoV-2 infected cell lines

Chimeric reads between human and SARS-CoV-2 RNA have been identified in RNA-seq data from infected cells in two recent studies (Zhang et al., 2020;Ying et al., 2021), presumed to be transcribed from reversed transcribed SARS-CoV-2 RNA integrated into the host DNA. To confirm these findings and exclude alternative origins of virus-host chimeric reads, we analysed public RNA-seq datasets of cells infected with unrelated RNA viruses or SARS-CoV-2, and lung samples from a COVID-19 patient and a healthy control, using a standard pipeline, also used in the previous studies (Zhang et al., 2020;Ying et al., 2021). As expected, reads mapping to the SARS-CoV-2 genome were readily found in samples infected with this virus (Figure 1A). A549 cells overexpressing *ACE2*, encoding the cellular receptor for SARS-CoV-2, from an adenoviral vector, and Calu3 cells showed the highest number of viral reads, with parental A549 cells and normal human bronchial epithelial (NHBE) cells showing lower read numbers (Figure 1A). Very few reads were identified in lung tissue samples taken from the COVID-19 patient and no SARS-CoV-2-mapping reads were identified in uninfected cell lines or those infected with unrelated viruses (Figure 1A). In agreement with earlier reports (Zhang et al., 2020;Ying et al., 2021), we identified host-viral junctions in SARS-CoV-2 infected cell lines, in direct proportion with the number of SARS-CoV-2 non-chimeric reads (Figure 1B). Supported human-SARS-CoV-2 chimeric reads constituted between 0.002% and 0.14% of all SARS-CoV-2-mapping reads found in infected cell lines, in line with the proportion of chimeric reads reported in earlier studies (Zhang et al., 2020;Ying et al., 2021), while no chimeric junctions were called in lung samples. Also in agreement with earlier studies, the viral parts of human-SARS-CoV-2 chimeric reads preferentially aligned to the 3’ end of the viral genome, mirroring general transcriptional activity of the viral genome (Figure 1C). Thus, human-SARS-CoV-2 chimeric reads are detectable in RNA-sed data, with the viral part donated more frequently from the highest expressed 3’ end of the viral genome.

**FIGURE 1.**
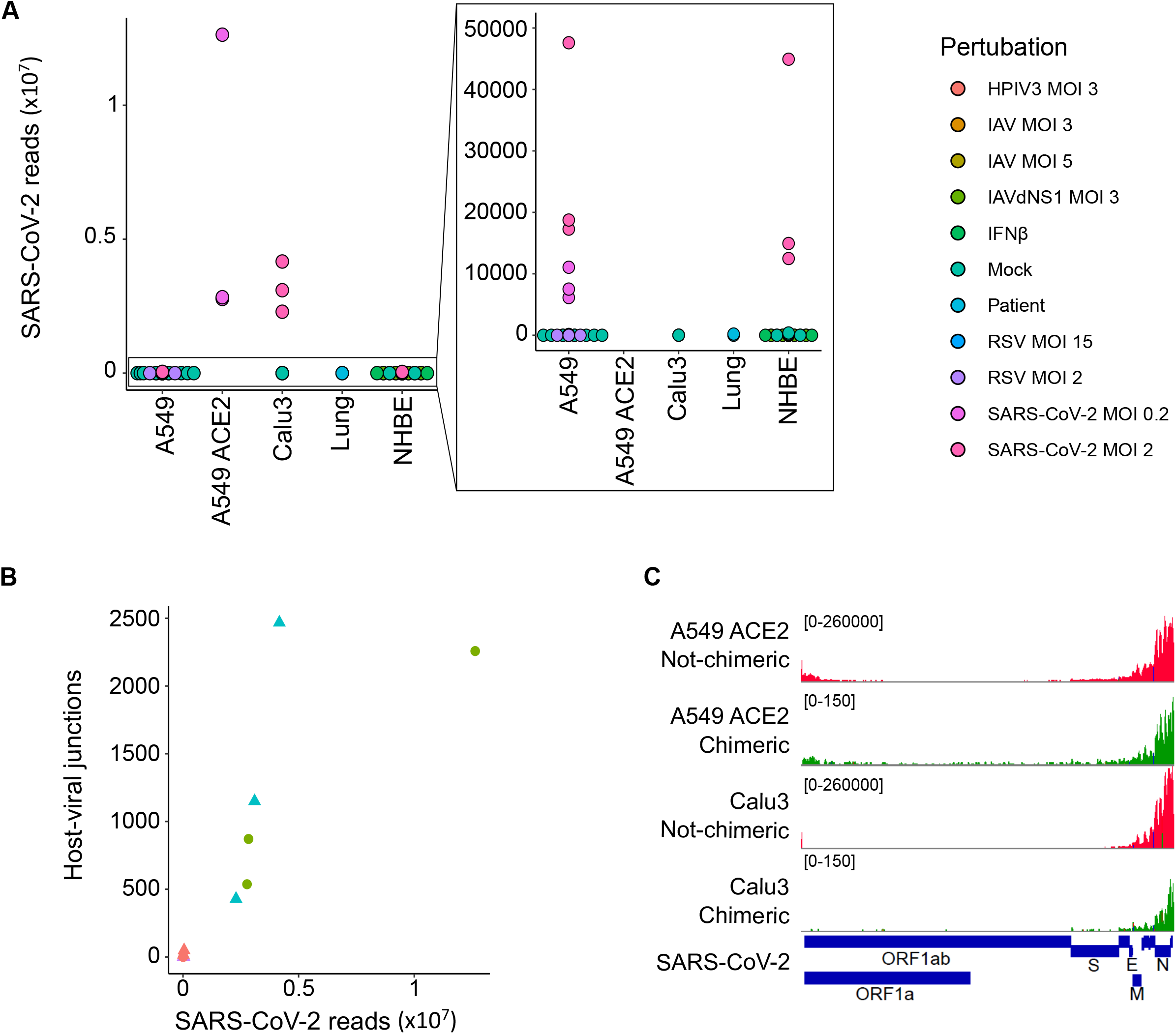
Detection of human-SARS-CoV-2 chimeric reads in RNA-sed data. **(A)** Number of non-chimeric reads uniquely aligning to SARS-CoV-2 genome in RNA-seq data (GSE147507) from parental A549 cells, A549 cells overexpressing ACE2 (A549 ACE2), Calu3 cells, NHBE cells, and lung tissue from a COVID-19 patient. The cells were infected or not (Mock) with human parainfluenza virus type 3 (HPIV3), influenza A virus (IAV), influenza A virus lacking the antiviral NS1 gene (IAVdNS1), respiratory syncytial virus (RSV) or SARS-CoV-2, at different multiplicities of infection (MOIs), or treated with recombinant IFNβ (IFNβ). **(B)** Number of human-SARS-CoV-2 junctions plotted against non-chimeric SARS-CoV-2-mapping reads in the same samples. **(C)** Alignment of human-SARS-CoV-2 chimeric and non-chimeric RNA-seq reads from SARS-CoV-2 infected A549 ACE2 and Calu3 cells across the SARS-CoV-2 genome, visualised on IGV.

### Non-canonical origin of the human part in human-SARS-CoV-2 chimeric reads

We next examined the possible location of the human sequence part found in human-SARS-CoV-2 chimeric reads along the human genome. Of all chimeric reads identified in SARS-CoV-2 infected A549 ACE2 cells, between 12.2% and 17.7% were formed between human mitochondrial and viral RNA (Figure 2A). In SARS-CoV-2 infected Calu3 cells mitochondrial RNA-SARS-CoV-2 chimeric reads comprised between 6.5% and 7.2% of total chimeric reads. Between 4.8% and 6.7% of chimeric reads in A549 ACE2 cells aligned to the ACE2 gene (Figure 2A). However, no ACE2-SARS-CoV-2 chimeric reads were found in other cell lines, including A549 cells. ACE2 overexpression in A549 ACE2 cells was achieved via transfection with an ACE2-expressing adenoviral vector (Blanco-Melo et al., 2020). As ACE2 in A549 ACE2 cells is transcribed from the adenoviral vector, chimeric ACE2-SARS-CoV-2 RNA-seq reads found in these cells would have required integration into the episomal adenoviral vector. Together, human-SARS-CoV-2 chimeric reads where the human part was donated by mitochondrial RNA or ACE2 RNA transcribed from the episomal adenoviral vector accounted for approximately a quarter of all chimeric reads (Figure 2A).

**Figure 2.**
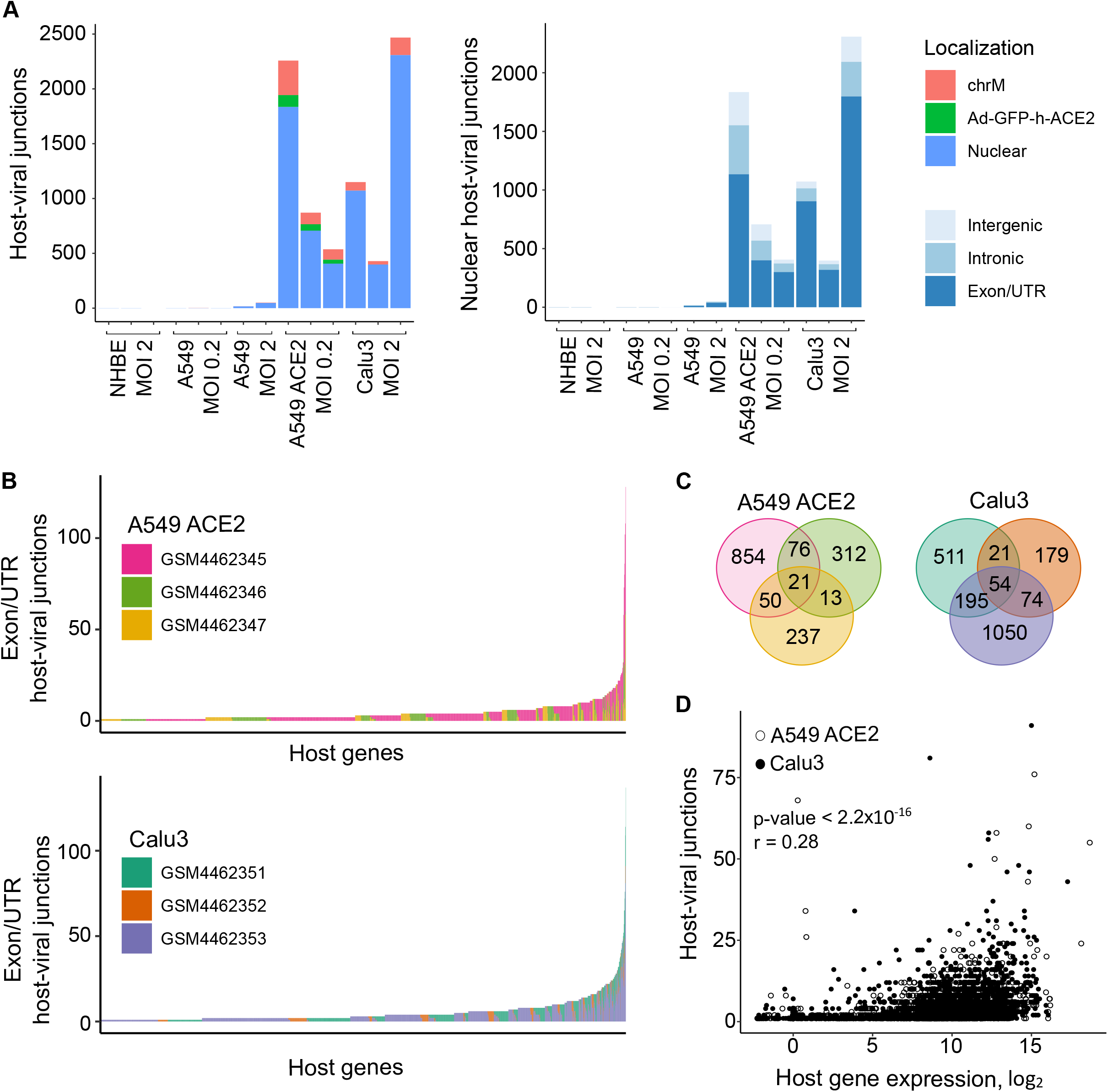
Characteristics of human sequence part of human-SARS-CoV-2 chimeric reads. **(A)** *Left*, number of chimeric junctions identified in the indicated samples, according to the origin of the human sequence part (chrM, mitochondrial DNA; Ad-GFP-h-ACE2, ACE2-encoding adenoviral vector episomal DNA, Nuclear, nuclear DNA). *Right*, number of chimeric junctions where the human sequence part aligns to nuclear human DNA, according to position relative to annotated genes and exons. Each bar presents each of the triplicate samples. **(B)** Number of chimeric junctions located within coding regions of individual human nuclear genes in A549 ACE2 and Calu3 RNA-seq datasets. Nuclear genes donating the human sequence part are plotted on x-axis and each bar represents an individual gene, colour-coded according to the triplicate sample in which it was found. **(C)** Overlap of host nuclear genes found in chimeric reads between the triplicate A549 ACE2 and Calu3 samples. **(D)** Correlation between the number of human-SARS-CoV-2 chimeric reads and the level of human donor gene expression.

The remaining human-SARS-CoV-2 chimeric reads aligned to nuclear genome. Of these, between 56.6% and 84.3% were located within annotated coding exons or untranslated regions (UTRs), whereas chimeric reads aligning to introns or intergenic regions were far fewer (Figure 2A). Notably, certain host genes contributed disproportionally to chimeric reads (Figure 2B). A549 ACE2 and Calu3 cells, 21 and 54 genes, respectively donated the human part of chimeric reads found in all three replicates of each cell line, and 139 and 290 genes, respectively contributed to chimeric reads in two of the replicates (Figure 2C). Host genes with higher contribution to chimeric reads also tended to be expressed at higher levels (Figure 2D). The recurrent contribution (between 14% and 45%) of the same highly expressed genes to chimeric reads in independent replicates of A549 ACE2 and Calu3 cell infection with SARS-CoV-2 indicates that the process that creates these chimeric reads was efficiently repeated in each replicate.

### Alternative mechanisms creating chimeric reads in RNA-seq libraries

In addition to reverse transcription and integration of viral RNA, followed transcription of the integrated copy, several alternative mechanisms might explain formation of chimeric RNA, such as genomic rearrangements, trans-splicing or transcriptional slippage (Yang et al., 2013). However, joining of transcripts from separate chromosomes or between host and viral RNA remains theoretical. An alternative mechanism for formation of inter-chromosomal chimeric reads in RNA-seq libraries has also been proposed (Li et al., 2009;Peng et al., 2015;Xie et al., 2016). This involves consecutive reverse transcription reactions, were cDNA sequences created during one reverse transcription reaction may prime reverse transcription of an unrelated RNA sequence through complementarity provided by small homologous sequences (SHS) (Li et al., 2009;Peng et al., 2015;Xie et al., 2016). The generation of artificial chimeric sequences via consecutive reverse transcription reactions is indirectly supported by the presence of mitochondrial RNA-nuclear RNA (mtRNA-nRNA) fusions in public expression sequence tags (ESTs) databases. The spatial separation of mitochondrial and nuclear DNAs negates transcriptional slippage or trans-splicing, leaving consecutive reverse transcription reactions through SHS-mediated priming as a possible cause.

To address the possibility that human-SARS-CoV-2 chimeric reads were formed via SHS-mediated priming during RNA-seq library construction, we first searched for mtRNA-nRNA chimeric reads, in order to assess the frequency of SHS at the junction of artifactual chimeric reads. Between 16% and 28% of analysed mtRNA-nRNA junctions exhibited an overlap of 3 or more nucleotides between mitochondrial and nuclear sequences (Figure 3A). We next looked for similar SHS across junctions of human-SARS-CoV-2 chimeric reads (Figure 3B, C). Between 14% and 16% of human-SARS-CoV-2 junctions had 3 or more overlapping nucleotides, which was comparable with their proportion in mtRNA-nRNA junctions (Figure 3A, B). Thus, SHS-mediated priming may be responsible for at least a fraction of human-SARS-CoV-2 chimeric reads detected in RNA-seq libraries.

**Figure 3.**
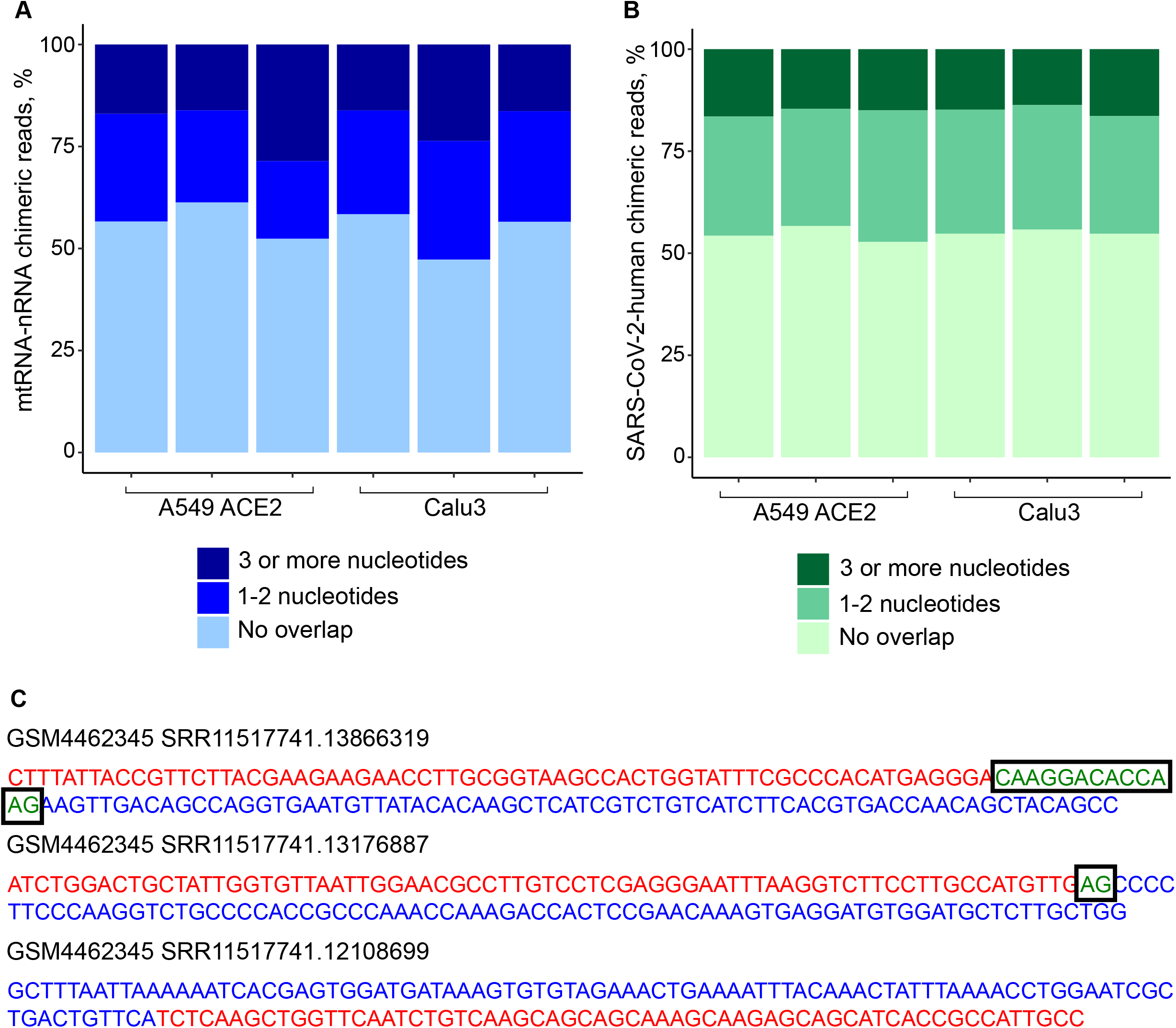
Sequence homology at the junctions of human-SARS-CoV-2 chimeric reads. **(A)** Sequence overlap between mitochondrial and nuclear DNA in mtRNA-nRNA chimeric reads. Each bar presents each of the triplicates of SARS-CoV-2 infected A549 ACE2 and Calu3 cells. **(B)** Sequence overlap between SARS-CoV-2 genomic RNA and the human genome in human-SARS-CoV-2 chimeric reads in the same samples. **(C)** Representative examples of human-SARS-CoV-2 chimeric reads with 13, 2 and 0 nucleotide overlap. SARS-CoV-2 and human genomic sequences are shown in red and blue letters, respectively. Overlapping sequences are shown in boxed green letters.

## DISCUSSION

The pandemic caused by SARS-CoV-2 that currently continues to spread globally (Hu et al., 2020), highlighted the need for deeper understanding of its interaction with the human host. The possible genomic integration of SARS-CoV-2 nucleic acids (Zhang et al., 2020;Ying et al., 2021) would have significant implications for host-viral interaction.

The somatic integration of a DNA copy of the RNA virus lymphocytic choriomeningitis virus (LCMV) in the murine host can provide a source of persistent antigen for the immune system (Klenerman et al., 1997). Similarly, persistence of somatically integrated SARS-CoV-2 DNA copies with coding potential could prolong presentation of viral antigens. However, analyses of intestinal biopsies several months after recovery from coronavirus disease 2019 (COVID-19), indicated the presence of SARS-CoV-2 RNA, as well as presumptive SARS-CoV-2 virions, consistent with on-going replication (Gaebler et al., 2021). Therefore, detection of persistent viral antigen may not necessarily indicate somatic SARS-CoV-2 integration.

Detection of chimeric reads between SARS-CoV-2 RNA and human RNA could also be indicative of somatic SARS-CoV-2 integration. Since detection of such chimeric reads in RNA-seq data would require transcription of the somatic integration, it would likely underestimate the total number of integrations. The high frequency of expressed somatic SARS-CoV-2 integrations reported (Zhang et al., 2020;Ying et al., 2021) was, therefore, unexpected. However, the majority of chimeric human-SARS-CoV-2 RNA reads may have a different origin. We identified chimeric reads between SARS-CoV-2 RNA and mitochondrial RNA, which were unlikely to have resulted from transcription of SARS-CoV-2 DNA copies integrated into mitochondrial DNA. If these reads were the result of SARS-CoV-2 integration into mitochondrial DNA, this would require mitochondrial import of viral cDNA and of components of canonical non-homologous end joining (NHEJ) process. While low levels of NHEJ had been reported in mitochondria, no evidence of viral DNA retrotransposition into the mitochondrial genome has yet been reported. Similarly, we identified chimeric reads between SARS-CoV-2 and RNA transcribed from the adenoviral vector used to overexpress *ACE2*, in target cells (Blanco-Melo et al., 2020), which would have necessitated integration of SARS-CoV-2 DNA copies in episomal adenoviral DNA. The finding that up to 24% of chimeric reads were formed between SARS-CoV-2 RNA and RNA transcribed from mitochondrial DNA or episomal adenoviral DNA suggested similarly artifactual generation of the remaining reads.

Chimeric reads between nuclear DNA-transcribed RNA and SARS-CoV-2 RNA involved host genes expressed at higher than average level. This correlation may have resulted from more probable detection of the higher expressed, than lower expressed genuine chimeric fragments. Alternatively, it could result from more frequent fortuitous joining, such as during RNA-seq library preparation for example, of SARS-CoV-2 RNA reads with the most abundant host gene RNA reads in the library. In support of the latter possibility, a substantial proportion of chimeric reads displayed complementarity, often over 10 nucleotides, in the joining region. Moreover, the substantially higher contribution of exonic than intronic or intergenic host sequences to human-SARS-CoV-2 chimeric reads is consistent with formation during RNA-seq library preparation, where exonic sequences are overrepresented relative to intronic or intergenic sequences. Together, these findings argue against frequent genomic integration and subsequent expression of SARS-CoV-2 RNA, as least at the level that can be determined by RNA-seq data analysis, and similar conclusions were reached by independent analysis (Yan et al., 2021).

## AUTHOR CONTRIBUTIONS

AK analysed the data. AK and GK wrote the manuscript.

## FUNDING

This work was supported by the Francis Crick Institute (FC001099), which receives its core funding from Cancer Research UK, the UK Medical Research Council and the Wellcome Trust; and by the Wellcome Trust (102898/B/13/Z). The funders had no role in study design, data collection and analysis, decision to publish, or preparation of the manuscript.

## ACKNOWLEDGMENT

The authors are grateful for assistance from the Scientific Computing Facility at the Francis Crick institute.

## Notes

### Competing Interest Statement

The authors have declared no competing interest.

## REFERENCES

Baertsch, R., Diekhans, M., Kent, W.J., Haussler, D., and Brosius, J. (2008). Retrocopy contributions to the evolution of the human genome. BMC Genomics 9, 466.

Blanco-Melo, D., Nilsson-Payant, B.E., Liu, W.C., Uhl, S., Hoagland, D., Møller, R., Jordan, T.X., Oishi, K., Panis, M., Sachs, D., Wang, T.T., Schwartz, R.E., Lim, J.K., Albrecht, R.A., and Tenoever, B.R. (2020). Imbalanced Host Response to SARS-CoV-2 Drives Development of COVID-19. Cell 181, 1036–1045.e1039.

Blinov, V.M., Zverev, V.V., Krasnov, G.S., Filatov, F.P., and Shargunov, A.V. (2017). Viral component of the human genome. Mol Biol 51, 205–215.

Bolger, A.M., Lohse, M., and Usadel, B. (2014). Trimmomatic: a flexible trimmer for Illumina sequence data. Bioinformatics 30, 2114–2120.

Burns, K.H., and Boeke, J.D. (2012). Human transposon tectonics. Cell 149, 740–752.

Buzdin, A.A. (2004). Retroelements and formation of chimeric retrogenes. Cell Mol Life Sci 61, 2046–2059.

Camacho, C., Coulouris, G., Avagyan, V., Ma, N., Papadopoulos, J., Bealer, K., and Madden, T.L. (2009). BLAST+: architecture and applications. BMC Bioinformatics 10, 421.

Coffin, J.M., Hughes, S.H., and Varmus, H.E. (1997). Retroviruses. Cold Spring Harbor (NY): Cold Spring Harbor Laboratory Press.

Dobin, A., Davis, C.A., Schlesinger, F., Drenkow, J., Zaleski, C., Jha, S., Batut, P., Chaisson, M., and Gingeras, T.R. (2013). STAR: ultrafast universal RNA-seq aligner. Bioinformatics 29, 15–21.

Esnault, C., Maestre, J., and Heidmann, T. (2000). Human LINE retrotransposons generate processed pseudogenes. Nat Genet 24, 363–367.

Feschotte, C., and Gilbert, C. (2012). Endogenous viruses: insights into viral evolution and impact on host biology. Nat. Rev. Genet 13, 283–296.

Gaebler, C., Wang, Z., Lorenzi, J.C.C., Muecksch, F., Finkin, S., Tokuyama, M., Cho, A., Jankovic, M., Schaefer-Babajew, D., Oliveira, T.Y., Cipolla, M., Viant, C., Barnes, C.O., Bram, Y., Breton, G., Hägglöf, T., Mendoza, P., Hurley, A., Turroja, M., Gordon, K., Millard, K.G., Ramos, V., Schmidt, F., Weisblum, Y., Jha, D., Tankelevich, M., Martinez-Delgado, G., Yee, J., Patel, R., Dizon, J., Unson-O’brien, C., Shimeliovich, I., Robbiani, D.F., Zhao, Z., Gazumyan, A., Schwartz, R.E., Hatziioannou, T., Bjorkman, P.J., Mehandru, S., Bieniasz, P.D., Caskey, M., and Nussenzweig, M.C. (2021). Evolution of antibody immunity to SARS-CoV-2. Nature.

Hu, B., Guo, H., Zhou, P., and Shi, Z.-L. (2020). Characteristics of SARS-CoV-2 and COVID-19. Nature Reviews Microbiology.

Klenerman, P., Hengartner, H., and Zinkernagel, R.M. (1997). A non-retroviral RNA virus persists in DNA form. Nature 390, 298–301.

Li, X., Zhao, L., Jiang, H., and Wang, W. (2009). Short homologous sequences are strongly associated with the generation of chimeric RNAs in eukaryotes. J Mol Evol 68, 56–65.

Liao, Y., Smyth, G.K., and Shi, W. (2014). featureCounts: an efficient general purpose program for assigning sequence reads to genomic features. Bioinformatics 30, 923–930.

Love, M.I., Huber, W., and Anders, S. (2014). Moderated estimation of fold change and dispersion for RNA-seq data with DESeq2. Genome Biol 15, 550.

Peng, Z., Yuan, C., Zellmer, L., Liu, S., Xu, N., and Liao, D.J. (2015). Hypothesis: Artifacts, Including Spurious Chimeric RNAs with a Short Homologous Sequence, Caused by Consecutive Reverse Transcriptions and Endogenous Random Primers. J Cancer 6, 555–567.

Richardson, S.R., Salvador-Palomeque, C., and Faulkner, G.J. (2014). Diversity through duplication: whole-genome sequencing reveals novel gene retrocopies in the human population. Bioessays 36, 475–481.

Robinson, J.T., Thorvaldsdóttir, H., Winckler, W., Guttman, M., Lander, E.S., Getz, G., and Mesirov, J.P. (2011). Integrative genomics viewer. Nat Biotechnol 29, 24–26.

Staszak, K., and Makałowska, I. (2021). Cancer, Retrogenes, and Evolution. Life (Basel) 11.

Xie, B., Yang, W., Ouyang, Y., Chen, L., Jiang, H., Liao, Y., and Liao, D.J. (2016). Two RNAs or DNAs May Artificially Fuse Together at a Short Homologous Sequence (SHS) during Reverse Transcription or Polymerase Chain Reactions, and Thus Reporting an SHS-Containing Chimeric RNA Requires Extra Caution. PLoS One 11, e0154855.

Yan, B., Chakravorty, S., Mirabelli, C., Wang, L., Trujillo-Ochoa, J.L., Chauss, D., Kumar, D., Lionakis, M.S., Olson, M.R., Wobus, C.E., Afzali, B., and Kazemian, M. (2021). Host-virus chimeric events in SARS-CoV2 infected cells are infrequent and artifactual. bioRxiv, 2021.2002.2017.431704.

Yang, W., Wu, J.M., Bi, A.D., Ou-Yang, Y.C., Shen, H.H., Chirn, G.W., Zhou, J.H., Weiss, E., Holman, E.P., and Liao, D.J. (2013). Possible formation of mitochondrial-RNA containing chimeric or trimeric RNA implies a post-transcriptional and post-splicing mechanism for RNA fusion. PLoS One 8, e77016.

Ying, Y., Xiao-Zhao, L., Ximiao, H., and Li-Quan, Z. (2021). Exogenous coronavirus interacts with endogenous retrotransposon in human cells. Research Square.

Zhang, L., Richards, A., Khalil, A., Wogram, E., Ma, H., Young, R.A., and Jaenisch, R. (2020). SARS-CoV-2 RNA reverse-transcribed and integrated into the human genome. bioRxiv, 2020.2012.2012.422516.

